# Latitude-specific urbanization effects on life history traits in the damselfly *Ischnura elegans*

**DOI:** 10.1101/2023.02.27.530143

**Authors:** Gemma Palomar, Guillaume Wos, Robby Stoks, Szymon Sniegula

## Abstract

1. Many species are currently adapting to cities at different latitudes. Adaptation to urbanization may require eco-evolutionary changes in response to temperature and invasive species that may differ between latitudes.
2. Here, we studied single and combined effects of increased temperatures and invasive alien predator presence on the phenotypic response of replicated urban and rural populations of the damselfly *Ischnura elegans* and contrasted these between central and high latitudes.
3. Larvae were exposed to temperature treatments (current [20 ºC], mild warming [24 ºC], and heat wave [28 ºC; for high latitude only]) crossed with the presence or absence of chemical cues released by the spiny-cheek crayfish (*Faxonius limosus*), only present at the central latitude. We measured treatment effects on larval development time, mass, and growth rate.
4. Urbanization type affected all life history traits, yet these responses were often dependent on latitude, temperature, and sex. Mild warming decreased mass in rural and increased growth rate in urban populations. The effects of urbanization on mass were latitude-dependent, with central-latitude populations having a greater phenotypic difference. Urbanization effects were sex-specific with urban males being lighter and having a lower growth rate than rural males. At the current temperature and mild warming, the predator cue reduced the growth rate, and this independently of urbanization level and latitude of origin. This pattern was reversed during a heat wave in high-latitude damselflies.
5. Our results highlight the context-dependency of evolutionary and plastic responses to urbanisation, and caution for generalizing how populations respond to cities based on populations at a single latitude.

## Introduction

Urbanization has emerged as a strong and widespread source of selection affecting plant and animal communities (Alberti, et al., 2017; Catullo et al., 2019). Urbanization is closely related to drivers of eco-evolutionary change such as temperature and invasive species. Within the cities, the dense concentration of pavement, buildings, and other surfaces that absorb and retain heat creates ‘urban heat islands’ in which temperature is higher (Tam et al., 2015). Urbanization may also favour the introduction of invasive species reducing diversity by competitive exclusion of native species (McKinney, 2006). These phenomena together with other factors related to urbanization, such as pollution or noise, create particular environments within the cities that drive evolutionary changes in organisms, e.g. at the phenotypic and physiological levels, and in their interactions with other organisms (Alberti, et al., 2017; Lambert et al., 2021).

Global warming and latitudial gradient might further interact wtih urbanization and alter organisms thermal plasticity and evolution (Verheyen et al. 2019). These plastic and evolutionary responses may differ across the species latitudinal distribution because populations from different latitudes encounter different seasonal time constraints (Stoks et al., 2012). Since plastic responses are considered the first response to buffer the effects of novel environmental stressors (Fox et al., 2019), investigating reaction norms of organisms from urban and rural populations at different latitudes in response to warming and stress caused by invasive alien predators may provide valuable insights to forecast species’ phenotypic response to the rapid local and global changes (Verheyen et al., 2019). Life history traits such as body size are responsive to urbanization, although the trends are taxon dependent. While a reduction in body size was observed in urban bumblebees (Eggenberger et al., 2019) and birds (Meillère et al., 2015), the opposite pattern was found in moths (Merckx, et al., 2018). Urbanization effects may also be sex-specific, e.g. urban males are bigger in butterflies (Kaiser et al., 2016). Available theories predict that when facing different ecological conditions, as generally observed along an urbanization or latitudinal gradient, organisms may be ranked from a slow- to fast-living continuum (‘pace-of-life’ syndrome; Brans & De Meester, 2018; Réale et al., 2010). This may be reflected by a shift towards a faster pace-of-life with accelerated growth, rapid development, and lower body mass as urbanization increases (Brans & De Meester, 2018; Debecker et al., 2016). Therefore, as urbanization is expected to affect life history strategies and patterns of covariation between traits, it is important to consider multiple life history traits, both sexes and other interacting drivers of evolutionary change such latitude-specific growth season length and warming, to better reflect anthropogenic impacts of cities on organisms.

In this study, we tested and contrasted the response of life history traits to single and combined thermal and invasive alien predator treatments between urban and rural populations from different latitudes. Special attention went to whether these life history responses were sex specific. We ran a common garden experiment on the damselfly *Ischnura elegans* from replicated rural and urban populations at high-(southern Sweden) and central- (southern Poland) latitudes. In damselflies, urban populations were shown to grow slower than rural populations at central latitudes (Tüzün & Stoks, 2021), but whether such pattern is consistent across the species latitudinal range requires further investigation. At the central latitude, the higher annual temperatures and associated longer growth season allow more generations per year, but less time per generation, thereby causing higher seasonal time constraints (Corbet et al., 2006; Stoks et al., 2012). As the effect of urbanization on life history may differ between mild warming and heat wave temperatures (e.g. 30 °C), we tested both in the same study. Besides a thermal treatment (mild warming and a heat wave), we imposed a treatment where we manipulated the presence of chemical predator cues from an invasive alien predator, the spiny-cheek crayfish, *Faxonius limosus*, which has been co-occurring with Polish *I. elegans* populations for several decades, but has not been yet reported in Sweden. The spread of invasive crayfish predator has been largely mediated by human activities and urban areas represent an attractive environment for this species (Reynolds & Souty-Grosset, 2011).

## Materials and Methods

### Study species

*Ischnura elegans* is a common damselfly in Europe (Dijkstra & Schröter, 2020). Across a latitudinal gradient, populations are characterized by different number of generations per year (voltinism). At central latitudes (including Poland), populations are generally uni- and bivoltine, meaning one or two generations per year, respectively (Corbet et al., 2006; Norling, 2021). At high latitudes (including Sweden), populations are usually uni- and semivoltine, meaning that one or two years are required for completing one generation, respectively.

### Study populations

Adult *I. elegans* females were collected from two urban and two rural ponds each in southern Sweden (hereafter, high latitude) and in southern Poland (hereafter, central latitude) (Fig. 1, Table S1) on 22-23 June 2021, as described in Sniegula et al. (2020). The distances between urban and rural sites ranged from 5 to 19 km at the high latitude and from 26 to 96 km at the central latitude. We quantified the level of urbanization based on the percentage of impervious surface from the high-resolution layer database (20 m resolution) (EEA, 2020). We created a circular buffer of 1 km around each sampling location and calculated the average value of imperviousness in each buffer using Quantum-GIS (QGIS Development Team, 2017). Ponds were defined as urban when the average percentage of impervious surface area within the buffer was above 20 % and as rural when the percentage was below 1.5 % (Brans & De Meester, 2018).

**Fig. 1.**
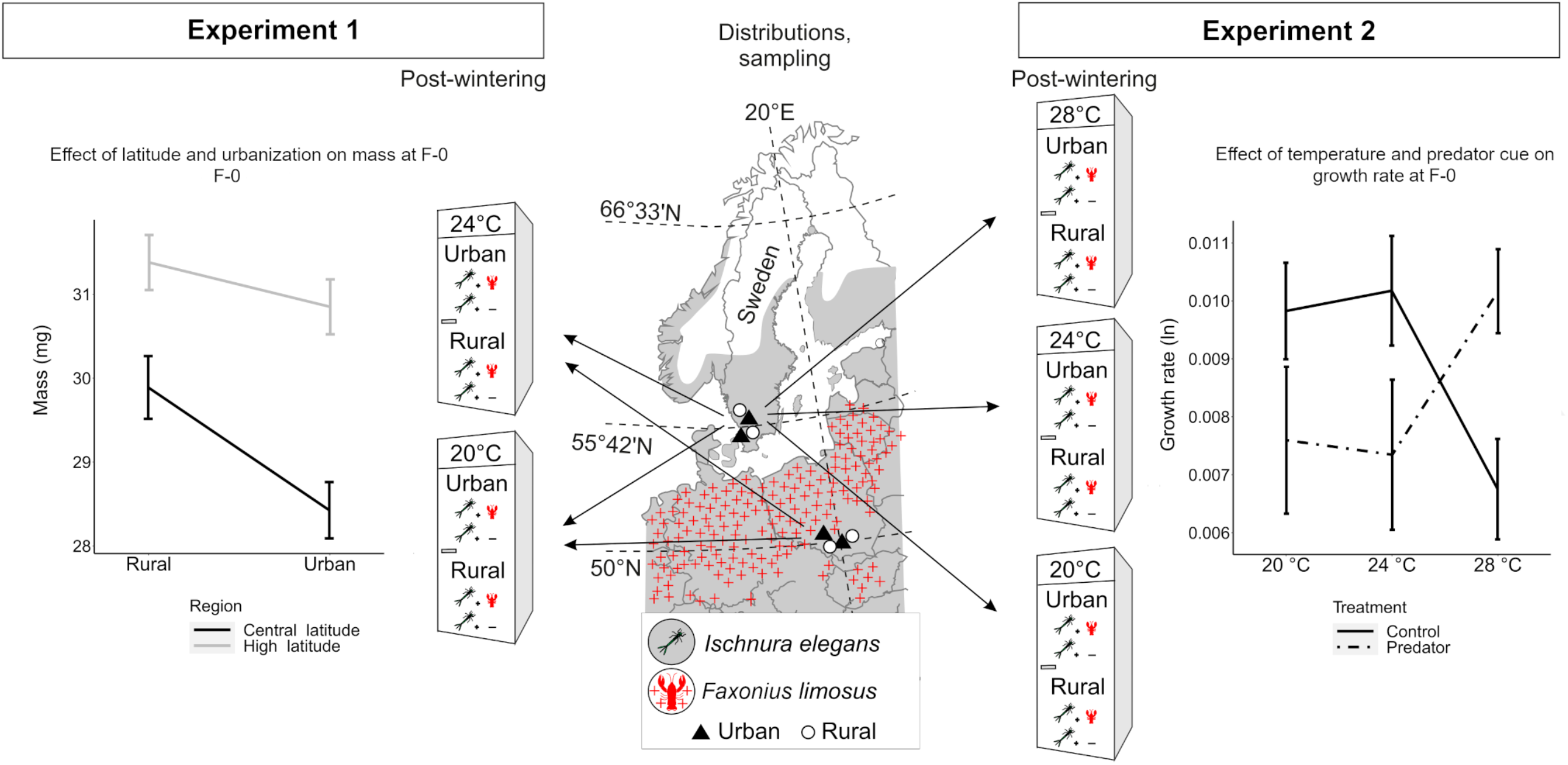
Summary of the experimental design with a map of our sampled populations in southern Sweden (high latitude) and southern Poland (central latitude). Geographic distribution of *Ischnura elegans* in central and northern Europe is shown in grey and occurrence of the spiny-cheek crayfish *Orconectes limosus* is depicted by red crosses. On the left side, we show the design of the experiment 1 focusing on rural and urban populations from central and high latitude reared at current (20 °C) and warming (24 °C) temperature. Illustrative plot shows the significant interaction latitude × urbanization for the final larval instar (F-0) mass. On the right side, we show the design of the experiment 2 focusing on urban and rural populations from high latitude reared at current (20 °C), warming (24 °C), and heat wave (28 °C) temperature and in a control or a predator cue treatment. Illustrative plot shows the significant interaction temperature × predator cue for the growth rate until F-0.

Water temperatures in the shallow parts of the collection ponds were estimated using the Lake Model FLake (2009) that closely matches the actual temperature measured in situ (Dinh Van et al., 2014). The modelled temperatures indicated minor differences in water temperature among ponds within- and between both latitudes (Fig. S1A and B). As the FLake model does not include impervious surface as a parameter and because water temperatures within a pond might vary depending on various parameters such as depth or sun exposure, we also placed temperature loggers (50 cm depth) in five ponds during summer and fall, and in one pond during an entire year (2021 and/or 2022; Fig. S1C and D). Based on FLake and dataloggers estimates, the average temperature in central- and high latitude ponds and between urban and rural ponds were similar and oscillated around 20 °C and were below 24

°C during summer months (except one record in one Polish pond, Fig. 1C). Based on this and on previous records of freshwater summer water temperatures at these latitudes (Debecker & Stoks, 2019; Dinh Van et al., 2014), we set the following experimental temperatures: 20 °C corresponding to the current mean summer water temperature, 24 ºC corresponding to the predicted increased temperature by 2100 under SSP8.5 scenario (Masson-Delmotte et al., 2021), and 28 ºC matching a simulated heat wave.

### The crayfish predator

The spiny-cheek crayfish *Faxonius limosus* is an invasive alien predator and an active colonizer that has locally co-occurred with the central-latitude *I. elegans* for at least 50 years but has not been reported in Scandinavia (Artportalen, 2022; Commission Implementing Regulation, 2016; Kouba et al., 2014). The spiny-cheek crayfish is widely reported in Poland in rural and urban areas (The General Directorate for Environmental Protection, 2018). The spread of the spiny-cheek crayfish is driven by human activities (aquaculture, trading), and natural spread in Europe (Reynolds & Souty-Grosset, 2011) and occurs frequently in warm temperatures (up to 25 ºC) and in polluted waters (Chucholl, 2016). Potential future invasions are more likely to occur in urban areas. Prior the experiment, *F. limosus* were collected from Kryspinów Lake in southern Poland (50°3′0.461′′N, 19°47′20.85′′E) and transported to the Institute of Nature Conservation of the Polish Academy of Sciences, Kraków, Poland (INC PAS). Three crayfishes were kept in an aquarium holding 50 L of dechlorinated tap water along with a control aquarium with 50 L of dechlorinated tap water. Crayfish collection and housing were done with permission from the Regional Directorate for Environmental Protection in Kraków (ref. OP.672.4.2021.GZ).

### Housing and temperature treatment

Adult female damselflies were individually placed in plastic cups with perforated lids and wet filter paper for egg laying. Females were kept at ca. 22°C and natural daylight (photoperiod). In total 80 clutches (10 clutches per population, hereafter families) were used in the experiment. The design of the experiment involved the same pre-winter and winter rearing conditions (hereafter, first part of the experiment) for all larvae. During post-winter rearing conditions (hereafter, second part of the experiment), larvae were exposed to different thermal treatments and, on the day of entering into the last instar prior emergence (F-0), larvae were exposed to the predator cue treatments for the next 5 days. The experiment ended at the end of this 5-day predator cue exposure period.

The first part of the experiment was carried out in plastic containers (size 22×16 cm, height 11 cm) filled with 1500 mL of dechlorinated tap water. Containers were kept in an incubator at 22 °C and a photoperiod of L:D 20:4h. In these containers, larvae were reared in groups. The photoperiod indicated the longest day length at the high latitude collection site, and was expected to create high development and growth rates in both studied latitudes, especially during post-winter conditions (Norling, 2021). Each container was supplied with a plastic structure to minimise predation among larvae. Since there were six different treatments after winter (i.e. during the second part of the experiment), larvae from each of the eight sampled populations were divided into six containers. At hatching, four larvae from each family were randomly placed in each of the six containers, totalling 40 larvae per container (4 larvae x 10 families). This approach ensured that each container started with exactly the same number of individuals from each family. Larvae were fed *ad libitum* with laboratory-cultured *Artemia nauplii*, twice a day on week days and once a day on weekend days. Furthermore, after three weeks of growing, we supplemented the feeding with live *Daphnia* sp. two times a week until autumn conditions. The position of the containers was randomized weekly within each incubator.

On 06 August 2021 (ca. four weeks after larvae had hatched), we started simulating autumn temperatures and photoperiods (hereafter, thermo-photoperiod) and, three weeks later, winter conditions. This procedure allowed larvae to experience a winter diapause, as it occurs in nature (Corbet 2006; Norling 2021). With a weekly interval, we gradually reduced the initial thermo-photoperiod from 22 °C and 20:4 h to 6 ºC and 0:24 h L:D (simulated winter conditions). A detailed description of rearing thermo-photoperiods during the entire experiment are presented in Fig. S2. During the simulated winter, larvae were fed three times a week with *Artemia nauplii*. On 22 November 2022, we started the second part of the rearing experiment. We transferred the larvae to individual 200 mL cups (height = 9 cm, diameter = 4 cm) filled with 100 mL of dechlorinated water and placed each cup into an incubator at 10 ºC and 4:20 h L:D. With a two-day interval, we gradually increased the thermo-photoperiod to the respective thermal treatment: 20 ºC (control), 24 ºC (mild warming) or 28 ºC (heat wave, high latitude only) with the same 20:4 h L:D photoperiod for all temperature treatments (Fig. S2). For logistic reasons we could not set the 28 °C temperature treatment group for the central-latitude populations. We therefore split our design into two experiments: Experiment 1 focusing on high- and central latitude populations raised at 20 °C and 24 °C, and Experiment 2 focusing on high-latitude populations raised at 20 °C, 24 °C and 28 °C.

Throughout the second part of the experiment, larvae were fed daily with *Artemia nauplii*. Because larvae came from different latitudes and because rearing temperature also affects larval development rate, larvae reached the F-0 stage at different dates. This led to larvae being exposed to the post-winter temperature treatment for different durations. For each larva, we reported the exact time in days at the temperature treatment during the second half of the experiment (‘thermal exposure duration’) and this time was included as a covariate in the statistical models (except for developmental time). In addition, when larvae entered F-0, we identified the sex of each individual to test for sex-specific responses to the treatments (Central latitude N _females_ = 73, N _males_ = 68; High latitude N _females_ = 158, N _males_ = 182).

### Predator cue treatment

When larvae entered the F-0 stage, we applied a five-day-long predator cue treatment. The water level in each cup was reduced to 67 mL and refilled with 33 mL of water from the crayfish aquarium (with predator cue) or the control aquarium (without predator cue). Cups were refilled every second day to keep the predator cue approximately constant, considering the length of predator cue biodegradation (Van Buskirk et al. 2014). Previous experiments have demonstrated that predator cues affect damselfly life history traits, also in case of short-time exposure (13 days exposure in Antoł & Sniegula, 2021; 3-9 days exposure in Van Dievel et al., 2016).

### Response variables

In total, 481 larvae survived and were phenotyped at the end of the experiment. Details on sample size for each treatment combination and response variable for both experiments are presented in Table S2. When larvae entered F-0 and before the application of the predator cue treatment, we quantified three traits: development time (DT; number of days between hatching and moult into F-0), wet mass (mass _F-0_), and growth rate until F-0 (GR _F-0_). Larval wet mass was measured with an electronic balance (Radwag AS.62) and GR _F-0_ was calculated as ln(mass _F-0_)/age _F-0_. After the five-day exposure to a predator cue, we measured the wet mass again (mass _final_) and calculated the growth rate over the five-day period: GR_final_ = [ln(mass _final_) – ln(mass _F-0_)]/5, as in McPeek et al. (2001) (Table S3).

### Statistical analyses

All analyses were performed in R (R Core Team, 2013; RStudio Team, 2015). For univariate statistics, we used generalized linear mixed-effects models (GLMMs). First, we ran a model selection analysis (MuMin r package; Barton & Barton, 2015) to select the most appropriate model for each phenotypic variable (DT, mass _F-0_, GR _F-0,_ and GR _final_). We included in the initial model the following predictors: latitude (only for experiment 1), sex, temperature, urbanization, predator (for GR _final_ only), and all the possible interactions; thermal exposure duration was added as a covariate (except for DT because this variable was strongly correlated with exposure duration) and population nested in latitude as a random factor. For each phenotypic variable, model selection analysis was based on the corrected Akaike’s information criteria for sample size (AICc) and weights as criteria to determine the best explanatory linear model by keeping only the most relevant predictors and interactions (Table S4). In Experiment 1, we compared urbanization type (rural vs. urban population), latitude (high- and central-latitude), temperature (20 ºC and 24 ºC), sex (males and females) and predator (for GR _final_ only). We ran specific models for each variable: DT _F-0_ with a Poisson distribution, and mass _F-0_, GR _F-0_ and GR _Final_ with Gaussian distributions, based on the output of the model selection analysis. The variables were normally distributed. In Experiment 2, we tested for effects of urbanisation type, temperature (20, 24 and 28°C) and predator (for GR _final_ only) in high-latitude populations. To fit the GLMMs, we used the function glmmTMB (glmmTMB package; Magnusson et al., 2017). P-values were obtained using the Wald chi-square test (Wald X²) implemented in the car package (Fox & Weisberg, 2015).

To analyse in detail the multivariate phenotypic responses of the larvae we applied Phenotypic Trajectory Analysis (PTA) following the procedure and R scripts described in Adams & Collyer (2009) and Collyer & Adams (2007). For this analysis, we used the measurements when larvae entered F-0 (hence, prior to the predator cue treatment). We used PTA to compare trajectories of high- and central-latitude populations in response to urbanization (evolutionary change expressed as a magnitude of phenotypic difference between rural and urban populations) and to temperature (plastic change expressed as the slope of the thermal reaction norm). We ran PTA both with the sexes pooled and with males and females separately. The procedure consists on a vector analysis in a multivariate space of phenotypic trait change. First, we conducted a PCA using DT _F-0_, mass _F-0,_ and GR _F-0_. To account for the fact that damselflies were exposed to the temperature treatment for different durations, we ran a GLM model for mass _F-0_ and GR _F-0_ with ‘thermal exposure duration’ as covariate and extracted the residuals that were subsequently used for the PCA and PTA analyses. Then, we created vectors connecting centroids of groups of individuals to calculate and compare the difference in length (magnitude) and angle (direction, θ) between groups. Significance in magnitude and direction were estimated using a permutation procedure (*N* = 1000 permutations); significant differences in the magnitude and direction between vectors were interpreted as different evolutionary or plastic trajectories.

## Results

### Experiment 1

#### Univariate response patterns

The results testing for the effects of the treatments on DT, mass, and GR are shown in Table 1. Urbanization had no main effect, but several interactions involving urbanization were significant. At 20 °C, urban larvae had a lower mass _F-0_ and GR _F-0_ compared to rural larvae. Mild warming (24 °C) reduced mass _F-0_, especially in rural larvae, and increased GR _F-0_, especially in urban larvae, eliminating the differences for these traits between urban and rural populations at 24 °C (interaction urbanization × temperature) (Fig. 2A and 2B). While urban and rural females did not differ in mass _F-0_ and GR _F-0_, urban males were lighter and showed a lower GR _F-0_ than rural males (interaction urbanization × sex) (Fig. 2C and 2D). Urbanization decreased larval mass, but in central-latitude individuals only (interaction latitude × urbanization) (Fig. 1); there was a larger difference in mass _F-0_ between latitudes in females than in males (interaction latitude × sex) (Fig. S3A). The interaction latitude × temperature was significant for GR _F-0_, with higher values of GR _F-0_ in central-compared to high latitude individuals but only at 24 °C (Fig. S3B). DT was not affected by urbanization type, but independently affected by latitude and temperature, with a shorter development time in central-latitude larvae and in larvae reared at 24 °C (Table S5; Fig. S4).

**Table 1.**
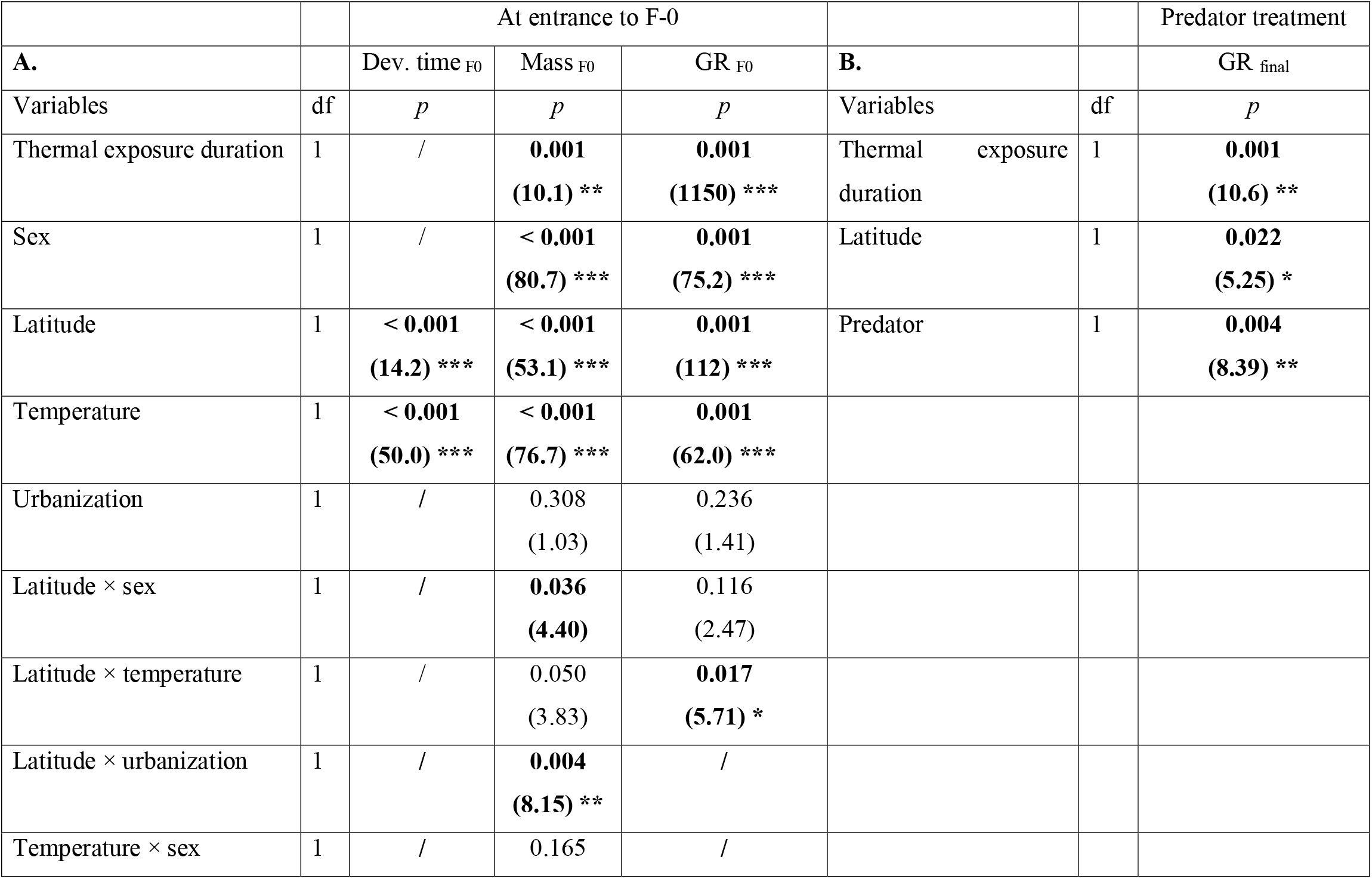
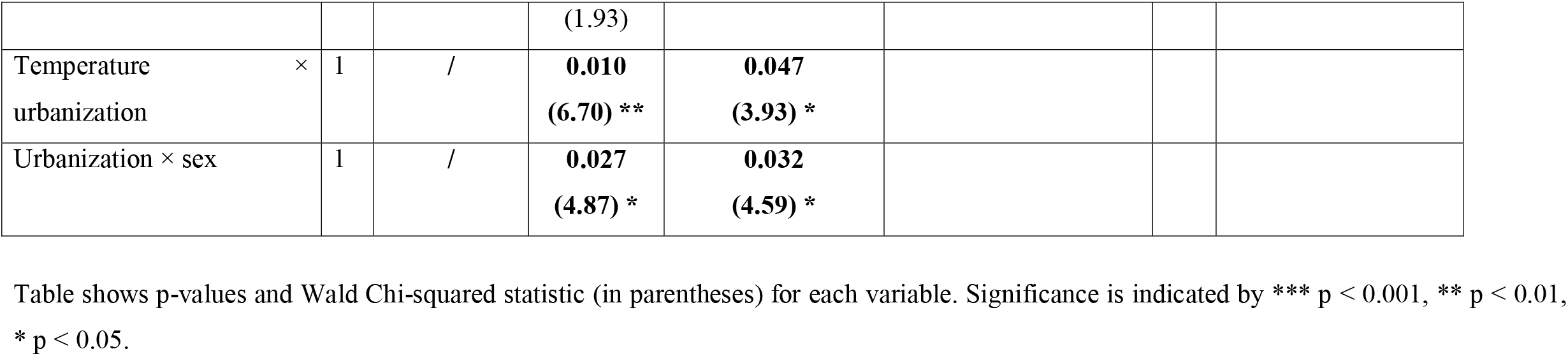
Results of the GLMM of the Experiment 1. Table shows effects of sex (females and males), latitude (central and high latitude), urbanization (rural and urban), temperature (20 °C and 24 °C) and of their interactions on larval development time, mass and growth rate (A) at entrance to F-0 (DT _F0_, Mass _F0_ and GR _F0_) and (B) during the five-days exposure period to predator cues (GR _final_). For each variable, we limited the analysis to the relevant predictors and interactions determined by the model selection analysis. The duration of the temperature treatment (post-winter ‘thermal exposure duration’) was included as covariate. Effects of random factors are not shown.

**Table 2.**
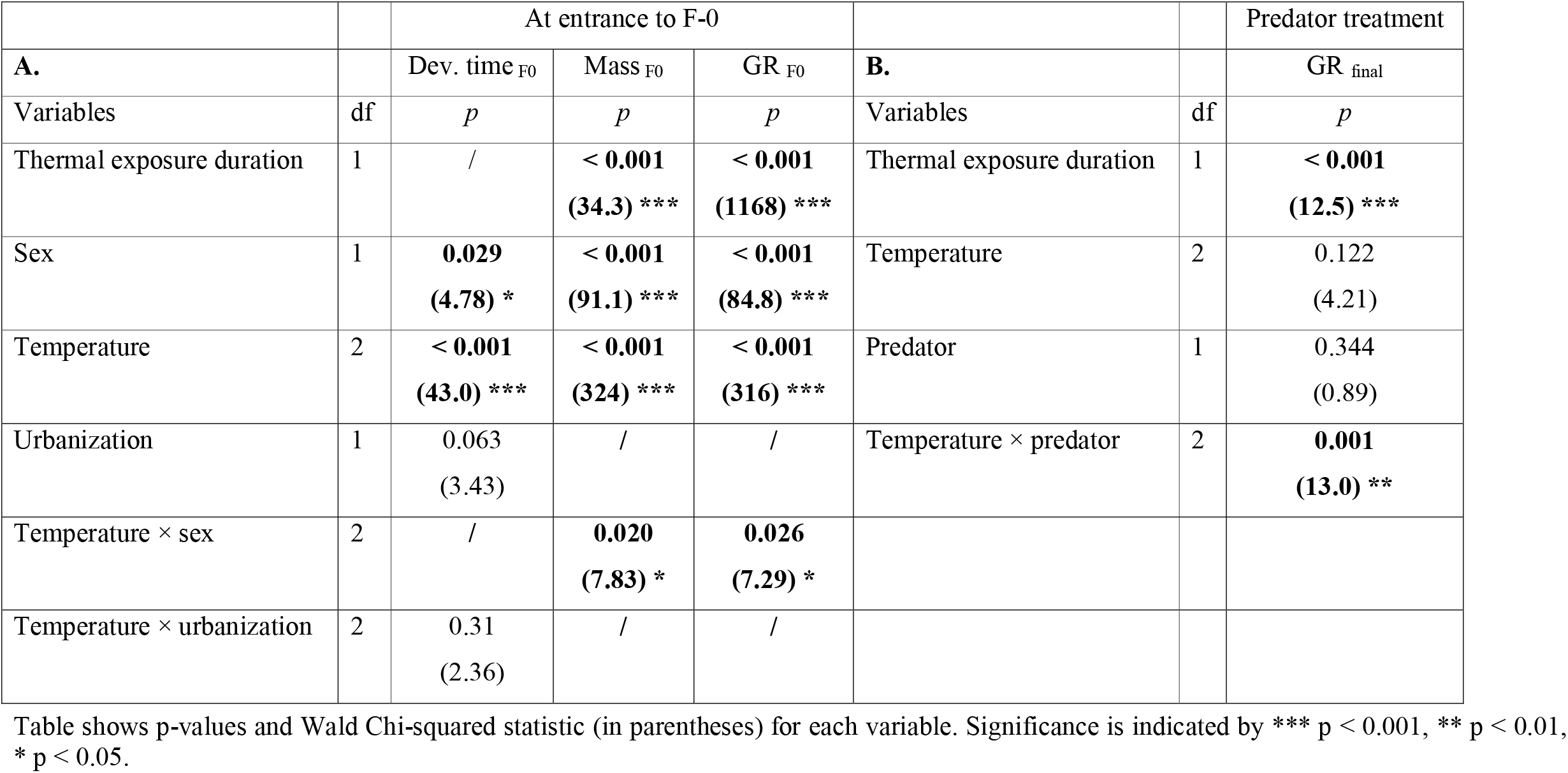
Results of the GLMM of the Experiment 2. Table shows effects of sex (females and males), urbanization (rural and urban), temperature (20 °C, 24 °C and 28 °C) and of their interactions on larval development time, mass and growth rate (A) at entrance to F-0 (Dev. time _F0_, Mass _F0_ and GR _F0_) and (B) during the five-days exposure period to predator cues (GR _final_). For each variable, we limited the analysis to the relevant predictors and interactions determined by the model selection analysis. The duration of the temperature treatment (post-winter ‘thermal exposure duration’) was included as covariate. Effects of random factors are not shown.

|  |  | At entrance to F-0 |  |  |  |  | Predator treatment |
| --- | --- | --- | --- | --- | --- | --- | --- |
| **A.** | | Dev. time $F_0$ | Mass $F_0$ | GR $F_0$ | **B.** | | GR $_{final}$ |
| Variables | df | *p* | *p* | *p* | Variables | df | *p* |
| Thermal exposure duration | 1 | / | **< 0.001**<br>(34.3) *** | **< 0.001**<br>(1168) *** | Thermal exposure duration | 1 | **< 0.001**<br>(12.5) *** |
| Sex | 1 | **0.029**<br>(4.78) * | **< 0.001**<br>(91.1) *** | **< 0.001**<br>(84.8) *** | Temperature | 2 | 0.122<br>(4.21) |
| Temperature | 2 | **< 0.001**<br>(43.0) *** | **< 0.001**<br>(324) *** | **< 0.001**<br>(316) *** | Predator | 1 | 0.344<br>(0.89) |
| Urbanization | 1 | 0.063<br>(3.43) | / | / | Temperature × predator | 2 | **0.001**<br>(13.0) ** |
| Temperature × sex | 2 | / | **0.020**<br>(7.83) * | **0.026**<br>(7.29) * |  |  |  |
| Temperature × urbanization | 2 | 0.31<br>(2.36) | / | / |  |  |  |

Table shows p-values and Wald Chi-squared statistic (in parentheses) for each variable. Significance is indicated by \*\*\* $p < 0.001$ , \*\* $p < 0.01$ , \* $p < 0.05$ .

**Fig 2.**
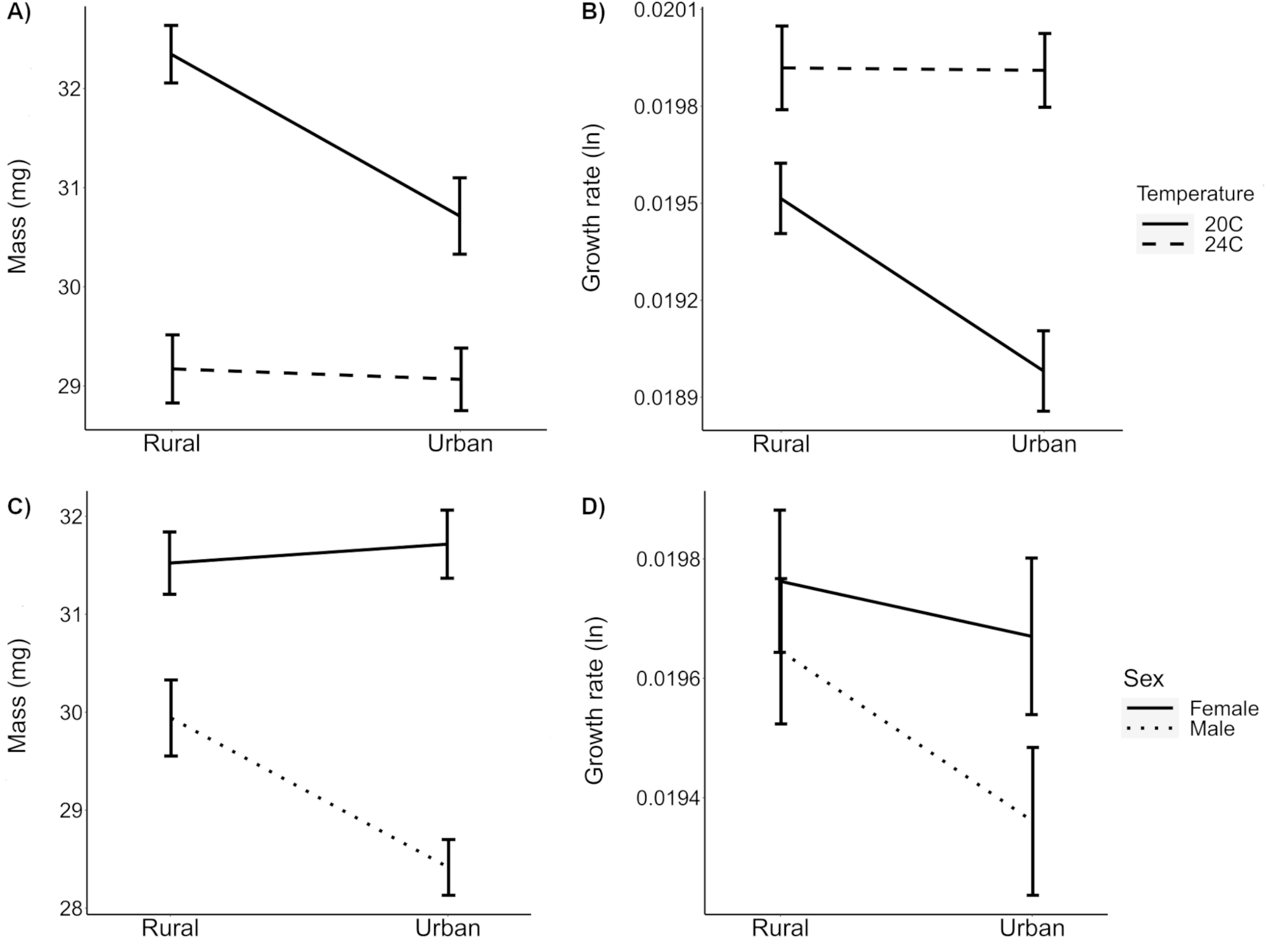
Larval (A) mass _F0_ and (B) growth rate _F0_ (GR _F0_) across urban and rural populations and current (20 °C) and warming (24 °C) temperature. Larval (C) mass _F0_ and (D) GR _F0_ across urban and rural populations for females and males in experiment 1.

During the five-day predator cue treatment, latitude and predator treatment had significant effect on GR _final_, with faster growth rate in central-latitude larvae and in the absence of predator cues (Table S5; Fig. S5). None of the interaction terms were significant for the GR _final._

#### Phenotypic Trajectory Analysis

For central-latitude populations, evolutionary changes in response to urbanization between larvae raised at 20 °C and 24 °C did not differ in length (Var_length_ = 0.39, *p* = 0.306) nor in direction (θ = 10.2°, *p* = 0.830). Evolutionary changes driven by urbanization pointed mostly to a lower mass and growth rate in urban areas (Fig. 3A). Plastic changes in response to mild warming temperature between rural and urban populations did not differ in length (Var_length_ = 0.27, *p* = 0.300) nor in direction (θ = 15.1°, *p* = 0.537). Urban and rural populations responded in a similar way to mild warming temperature with a shorter development time when temperature increased (Fig. 3C).

**Fig 3.**
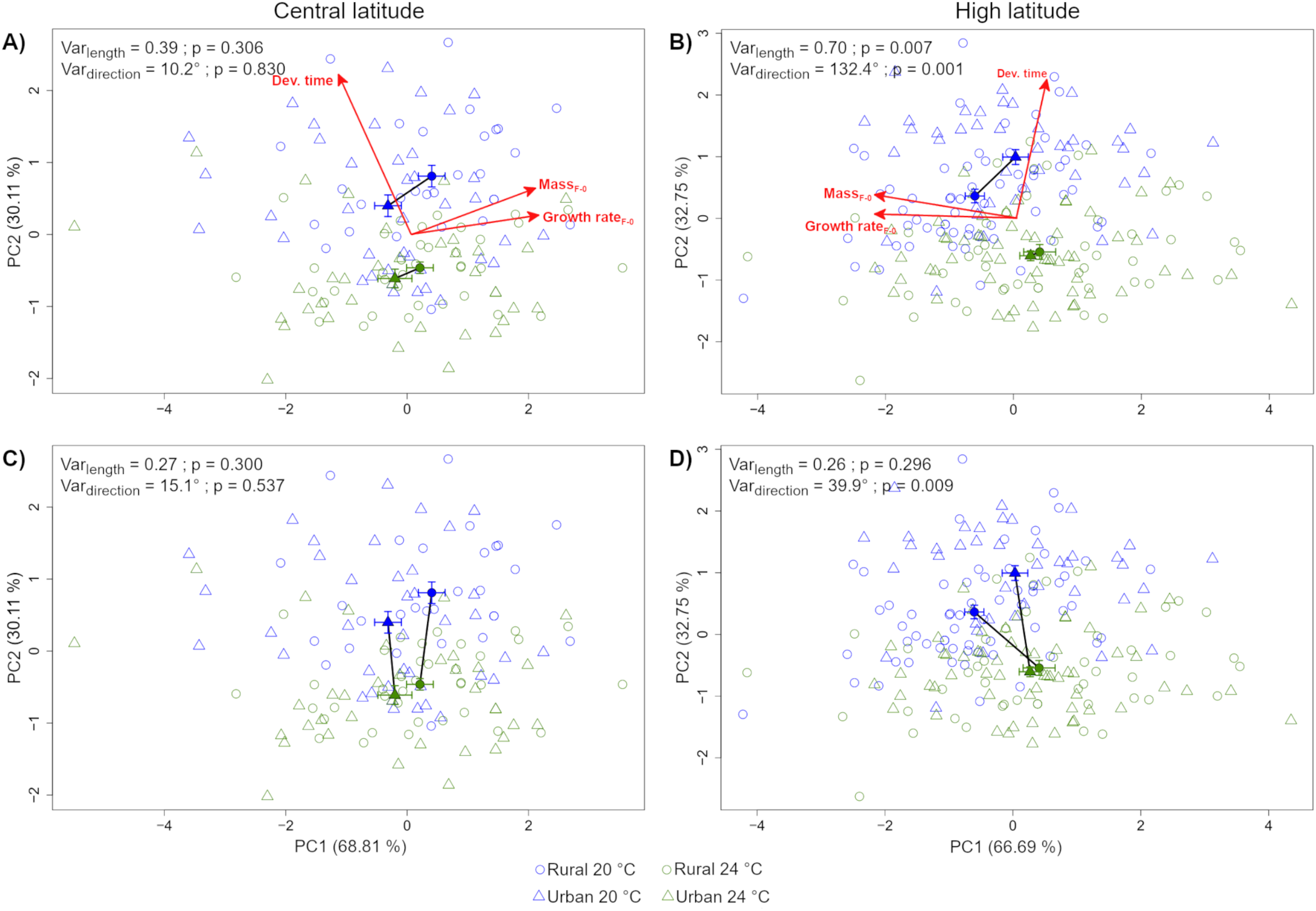
Principal component analysis showing evolutionary changes before the treatment with predator cue in response to urbanization at current (20 °C) and warming (24 °C) for (A) central- and (B) high latitude populations and plastic changes in response to temperature in rural and urban populations for (C) central- and (D) high latitude populations (sexes pooled). Rural and urban individuals are depicted by open circles and triangles respectively; temperature by colours (blue = 20 °C and green = 24 °C); filled circles and triangles correspond to the centroid of each group; solid lines connecting filled symbols represent the vector.

Next, we ran similar analyses for males and females separately. For males, we found the same patterns for evolutionary and plastic changes as with the sexes pooled (Fig. S6A and C). For females, we found different evolutionary trajectories (differences in direction only) between females raised at 20 °C and 24 °C (Fig. S7A); at 20 °C evolutionary changes in response to urbanization pointed to a decrease in mass and GR _F-0_ and the pattern was reversed at 24 °C. Thermal plasticity did not differ between rural and urban females (Fig. S7C).

For high-latitude populations, when sexes were pooled, evolutionary changes in response to urbanization differed both in length (Var_length_ = 0.70, *p* = 0.007) and direction (θ = 132.4°, *p* = 0.001) between the two temperature treatments (Fig. 3B). Urban larvae had a longer DT, and lower mass and GR than rural larvae at 20 °C, whereas there was little difference between urban and rural damselflies at 24 °C. Plastic changes in response to an increase in temperature showed no difference in magnitude between rural and urban populations (Var_length_ = 0.26, *p* = 0.296) but significant differences in direction (θ = 39.9°, p = 0.009) with a decrease in mass and GR at 24 °C only being present in rural populations (Fig. 3D).

When looking at each sex separately, for males, we found the exact same patterns for evolutionary (Fig. S6B) and plastic (Fig. S6D) changes as with the sexes pooled. For females, evolutionary changes differed only in direction between the temperature treatments; at 20 °C evolutionary changes pointed to an increase in development time in urban area and at 24 °C to an increase in mass and GR _F-0_ (Fig S7B). Plastic changes to mild warming temperature for females followed the same pattern as with the sexes pooled (Fig. S7D).

### Experiment 2

#### Univariate response patterns

In the set of high-latitude urban and rural larvae tested at the three temperatures, the score life-history traits (mass _F0_, DT _F0,_ and GR _F0_) were not affected by urbanization, but were affected by temperature and sex (Table S4). The interaction temperature × sex was significant for both mass _F-0_ and GR _F-0_, with higher values in mass _F-0_ and GR _F-0_ in females, especially at 24 °C (Fig. S8). GR _final_ was not affected by urbanization. Exposure to the predator cue during the five-day-long treatment reduced GR _final_ at 20°C and 24°C, but instead increased GR _final_ at 28°C (predator × temperature) (Fig. 1).

#### Phenotypic Trajectory Analysis

Phenotypic trajectory analysis showed that high-latitude populations exhibited different evolutionary changes in response to urbanization across the three temperature treatments (sexes pooled). We did not find significant variation in magnitude between the three vectors (Var_length_ = 0.02, *p* = 0.218) but we found significant differences in their direction (Var_direction_ = 3547.0, *p* = 0.001) (Fig. 4A). At 20 °C and 28°C, urbanization is mostly accompanied by longer development and lower mass _F-0_ and GR _F-0_, whereas at 24 °C differences between rural and urban individuals were rather small. For plastic changes, PTA revealed significant differences in length (Var_length_ = 0.720, *p* = 0.016) but not in direction (Var_direction_ = 10.0, *p* = 0.222) between the two vectors indicating a greater thermal plasticity in urban than in rural populations (Fig. 4B).

**Fig 4.**
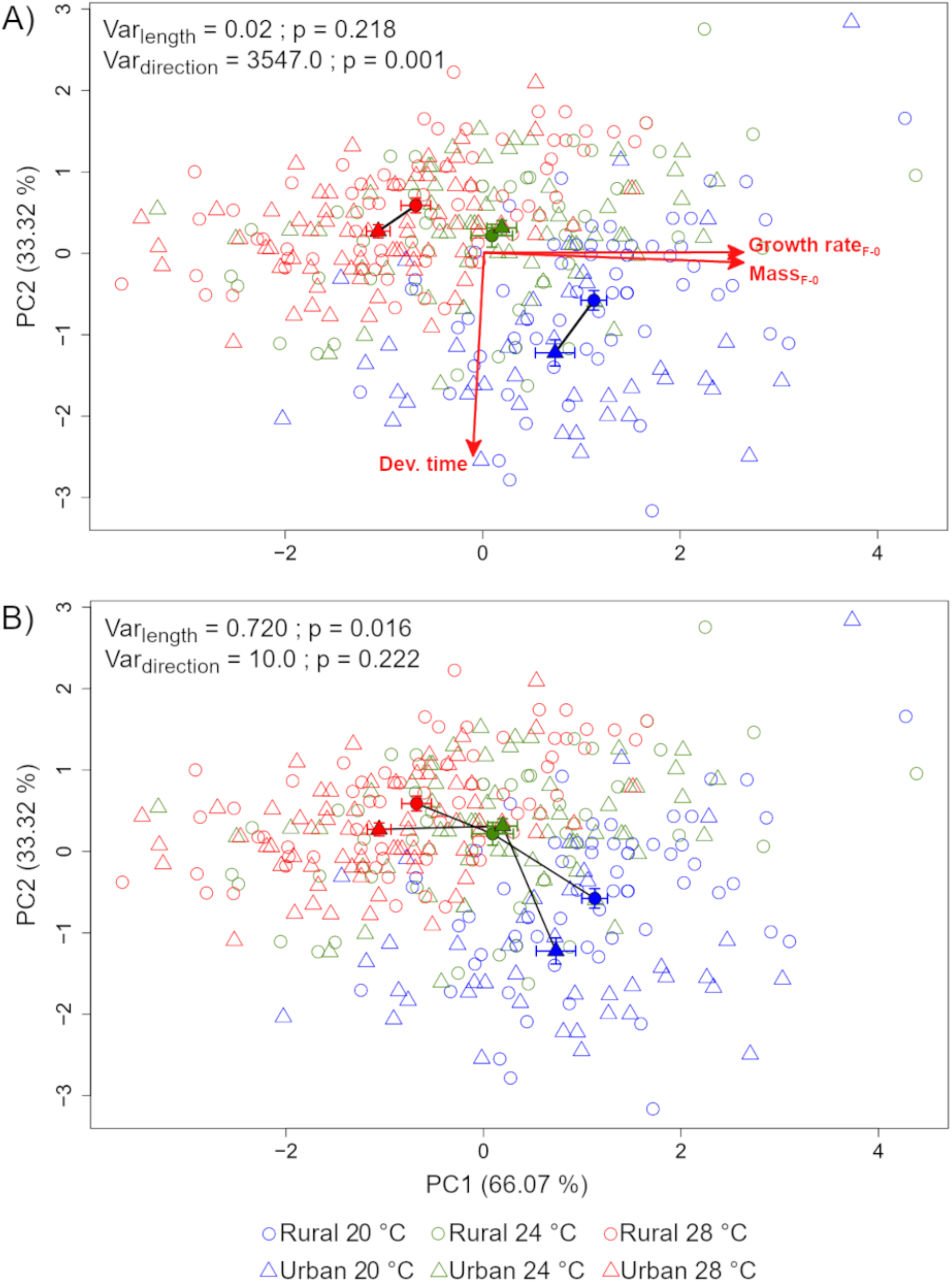
Phenotypic plasticity trajectory in response to (A) urbanization at current (20 °C), warming temperature (24 °C) and heat wave (28 °C) and to (B) temperature for rural and urban populations. Rural and urban individuals are depicted by open circles and triangles respectively; temperature by colours (blue = 20 °C, green = 24 °C and red = 28 °C); filled circles and triangles correspond to the centroid of each group; solid lines connecting filled symbols represent the vector.

When looking at each sex separately, evolutionary changes in response to urbanization were similar for males and females and followed the same patterns as with the sexes pooled (Fig. S9A and B). For plastic changes, no difference in both length and direction was found in males (Fig. S9C), whereas differences in length were found in females (Fig. S9D); with a greater thermal plastic response in females from urban than rural populations.

## Discussion

Here, we studied phenotypic adaptation in an ectotherm considering several life-history traits and interacting drivers of evolutionary change. Our results pointed to effects of urbanization in combination with temperature on mass and GR. Notably, the direction and magnitude of the evolutionary and plastic responses to temperature in life history traits (i.e. body mass and GR) were not uniform across studied latitudes. The response to a predator was rather independent of the latitudinal origin of the populations and of the urbanization type. Our results shed some light on the effect of interacting anthropogenic factors in ectotherms life-history traits and how these patterns are latitude dependent.

### Effects on individual traits

In Experiment 1, we found negative effects of urbanization on larval growth rate and mass, but these effects differed between latitudes and depended on temperature. This partially confirmed that urbanization can decrease mass in insects (Merckx, et al., 2018), and that mass decrease is temperature-dependent (Diamond et al., 2014). A mild warming temperature was sufficient to remove the differences between rural and urban larval mass and growth rate. Intriguingly, these two traits showed higher thermal plasticity at different urbanization types. While mass showed steeper thermal reaction norm in rural populations, urban populations were more plastic for GR. We might hypothesise that the effect of temperature was stronger than the effect of urbanization and that growth rate reached some physiological limits at 24 °C (growth rate did not increase at 28 °C in high latitude populations), however, the pattern was less clear for the mass.

Notably, the effects of urbanization were also latitude-dependent. The latitudinal difference may be attributed to different life-history strategies related to voltinism and seasonal time constraints. At central latitudes, the species produces, on average, one generation per year more than at high latitudes (Corbet et al., 2006; Norling, 2021). Therefore, central-latitude larvae are likely more time constrained: they develop within a shorter time, grow faster and reach lower mass than at high latitudes (as seen for the species between low and high latitudes, e.g. Debecker and Stoks 2019). A higher growth rate indeed often comes at a cost of having a smaller mass at emergence (Sniegula et al., 2018). In current study, latitudinal differences in mass seemed to be amplified by urbanization. Indeed, with climate change, there might be a further increase in time constraints in urban areas for central-latitude individuals that might arose from the heat-island effects (Chick et al., 2019). However, based on our field measurements, contemporary water temperatures in urban and rural ponds differed slightly (Fig. S1), and it is difficult to predict whether climate change will increase this difference in temperature.

Independently of urbanization and latitude, the predator cue negatively affected larval GR. In insects, exposure to a predator alters metabolism (Cinel et al., 2020) and foraging activity (Kohler & McPeek, 1989), with downstream effects on larval development and growth rate. Our results indicate that both central- and high-latitude populations reacted in a similar way to the predator cue in the traits we measured. These results contrast with a recent study showing a differential response in development time in *I. elegans* eggs when exposed to cues from non-native invasive predators (spiny-cheek and signal crayfish *Pacifastacus leniusculus*) (Antoł & Sniegula, 2021). However, an exposure to a phylogenetically related predator species, i.e. noble crayfish (*Astacus astacus*) present at both latitudes (Kouba et al., 2014), might enable predator cue recognition and trigger similar responses between the two latitudes (Anton et al., 2020).

Results from Experiment 2 did not provide evidence that urban and rural populations coped differently with a simulated heat wave, as previously demonstrated in damselflies (Tüzün & Stoks, 2021) and other ectotherms (Brans et al., 2017; Campbell-Staton et al., 2020). This may stem from the minor mean temperature differences between urban and rural ponds at our study sites or different experimental approaches. Yet, we found a significant effect of temperature in combination with predator cues. Damselflies grew faster in the absence of a predator cue, but only in current and warming temperature. Interestingly, exposure to the predator cues increased larval growth under a heat wave treatment. A combination of stressors may change a strategy in potential prey to escape predator exposure (Warkentin, 2011). Here, an accelerated larval growth in the presence of a predator cue under heat wave may be part of an escape strategy to reduce the time of exposure to predators, as previously shown in odonate species (Antoł & Sniegula, 2021; Stoks et al., 2012) and other taxa (Chivers et al., 2001).

In both experiments, we found multiple sex-specific effects on mass and GR with greater phenotypic differences between urbanization types and temperatures in males. These results supported previous studies in which sex-specific effects were found in ectotherms coping with various stressors, i.e. urbanization (Kaiser et al., 2016) and heat stress (Sniegula et al., 2017). Sex-specific effects are generally more pronounced in species with strong sexual dimorphism which is the case in damselflies, with females being usually larger and heavier than males (Corbet, 1999). Sex-specific effects are also common in protandrous species with a strong selection acting on males to emerge before females in order to maximise their mating opportunities (Badyaev, 2002). Hence, different selective pressures resulting in different life-history strategies and associated resource allocation patterns between males and females may lead to different types of trade-offs. For instance, female butterflies maintained a relative high body mass under different thermal conditions compared to males because mass was more important for females in reproduction than for males (Fischer & Fiedler, 2000). This is likely to be the case in odonates (Sokolovska et al., 2000) and matched our observations of mass of females being less affected by urbanization and temperature than males (at least until 24 °C; Fig. S3A).

### Multivariate approach

Our results indicate multivariate evolutionary change associated to urbanization at the high latitude since larvae from urban and rural sites differed in life history traits and in their thermal plasticity. In contrast, central-latitude damselflies exhibited similar direction and magnitude of evolutionary trajectories in response to urbanization in the two temperature treatments (Experiment 1). Similar results were found in high-latitude individuals in response to urbanization and additional simulation of heat wave (Experiment 2). The ‘pace-of-life’ syndrome predicts a shift towards a fast-living strategy when urbanization increases or towards lower latitudes, which was supported empirically in birds and invertebrates (Brans & De Meester, 2018; Charmantier et al., 2017; Debecker et al., 2016). We observed that only high latitude damselflies expressed a consistent decrease in mass and growth rate associated with urbanization at 20 ºC, which was further accompanied with an increase of development time. Hence, we found no support for a faster pace-of-life in the studied populations. One explanation might be the minor difference in water temperatures recorded in ponds from which rural and urban *I. elegans* were collected. However, in the damselfly *Coenagrion puella* a lower growth rate in urban compared to rural populations was also shown, despite water temperatures being up to 3.5 °C higher in urban than in rural ponds (Tüzün & Stoks, 2021). In that study, the population-specific growth pattern was explained by relatively lower temperature and hence shorter growing seasons in rural populations, i.e. compensation to time constraints by increased growth rate. Our results might, therefore, reflect selection caused by other sources of disturbance associated with the urban environment, e.g. different concentrations of pollutants. Notably, we demonstrated sex-specific evolutionary trajectories to urbanization at both latitudes. Therefore, the population origin (latitude) and the sex, which entail different life-history strategies or trade-offs, were more likely to trigger different trajectories than a shared response to urbanization.

Increased levels of environmental disturbance, including stress linked to suboptimal temperatures, in cities is expected to favour phenotypic plasticity (Alberti, et al., 2017). In contrast, rural and urban populations from the central latitude responded with similar magnitude and direction to mild warming by decreasing developmental time, whereas high-latitude populations showed similar plasticity in this trait to mild warming in terms of magnitude, but not in terms of direction. However, in response to the heat wave, we did find a greater magnitude of the plastic response in urban populations for GR _F-0_ and mass, matching the expectation of an increased phenotypic plasticity in urban populations. Different effects of urbanization on plasticity between the two latitudes may be partially caused by different life history strategies: lower latitude organisms tend to have shorter generation times (Corbet et al., 2006). Based on these results, we may infer that not only ‘urban heat island’ effects impose differential selective pressures on organisms across urbanisation gradients (Shochat et al., 2006), but also geographic origin and temperature exposition in rural and urban populations have the potential to adjust the damselfly phenotype.

## Conclusion

Despite the accumulating evidence that urbanization shapes phenotypes, and the expectation these evolutionary and plastic responses may differ among latitudes, the latter has rarely been tested. We showed that the damselfly responses to urbanization differed between latitudes. Notably, the latitude-specific responses to urbanisation were temperature-dependent, which could be explained by differences in life-history strategies across latitudes. Both urban and rural populations had the potential to produce a plastic response to warming, yet the magnitude and direction of the plastic changes differed between latitudes. Moreover, we added to the knowledge that the response to urbanization can differ between sexes. In contrast, urban and rural damselflies responded similarly to the presence of a predator cue, the later interacting with different temperatures. Our results highlight the context-dependency of evolutionary and plastic responses to urbanisation, and caution for generalizing of how populations respond to cities based on populations at a single latitude.

## Acknowledgments

We thank Andrzej Antoł, Maja Przybycień, Ola Rydzek and Antoni Żygadło that helped during the experiments. We thank Ulf Norling for valuable discussions on the topic. The research leading to these results was funded by the Norwegian Financial Mechanism 2014– 2021, project no. 2019/34/H/NZ8/00683 (ECOPOND). S.S. was further supported by the National Science Centre, Poland (project no. 2019/33/B/NZ8/00521) and the Institute of Nature Conservation Polish Academy of Sciences.

## Author contributions

GP, RS and SS conceived and designed the experiments. GP and SS performed the experiments. GW analysed the data. GW and SS led the writing of the manuscript; all authors contributed critically to the drafts and gave final approval for publication.

## Date availability statement

All data generated or analysed during this study are included in this article (and its supplementary information files).

