## Supplementary figures and images for "Latitude-specific urbanization effects on life history traits in the damselfly *Ischnura elegans*"

### Fig. S1.pdf

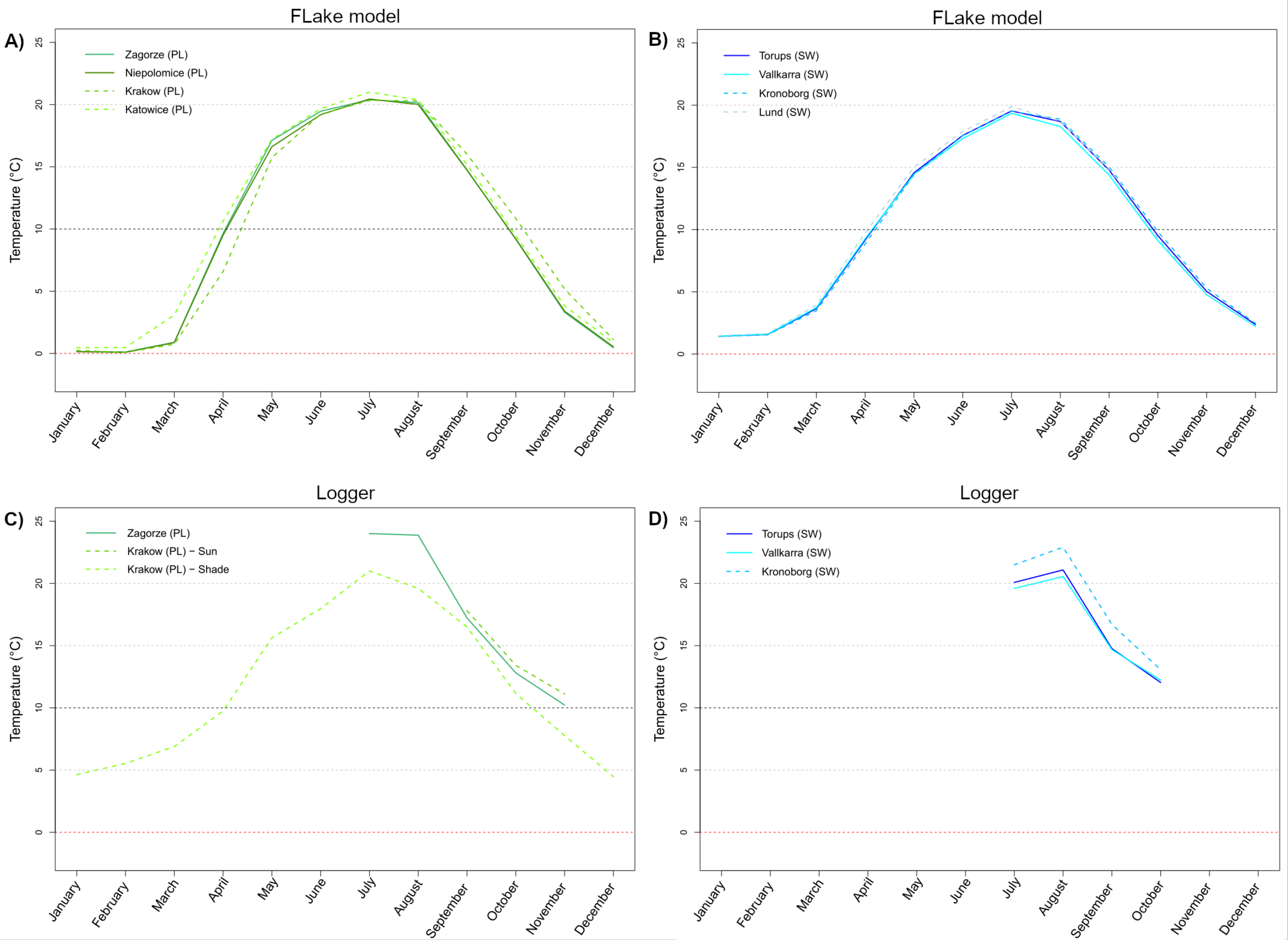

### Fig. S6.pdf

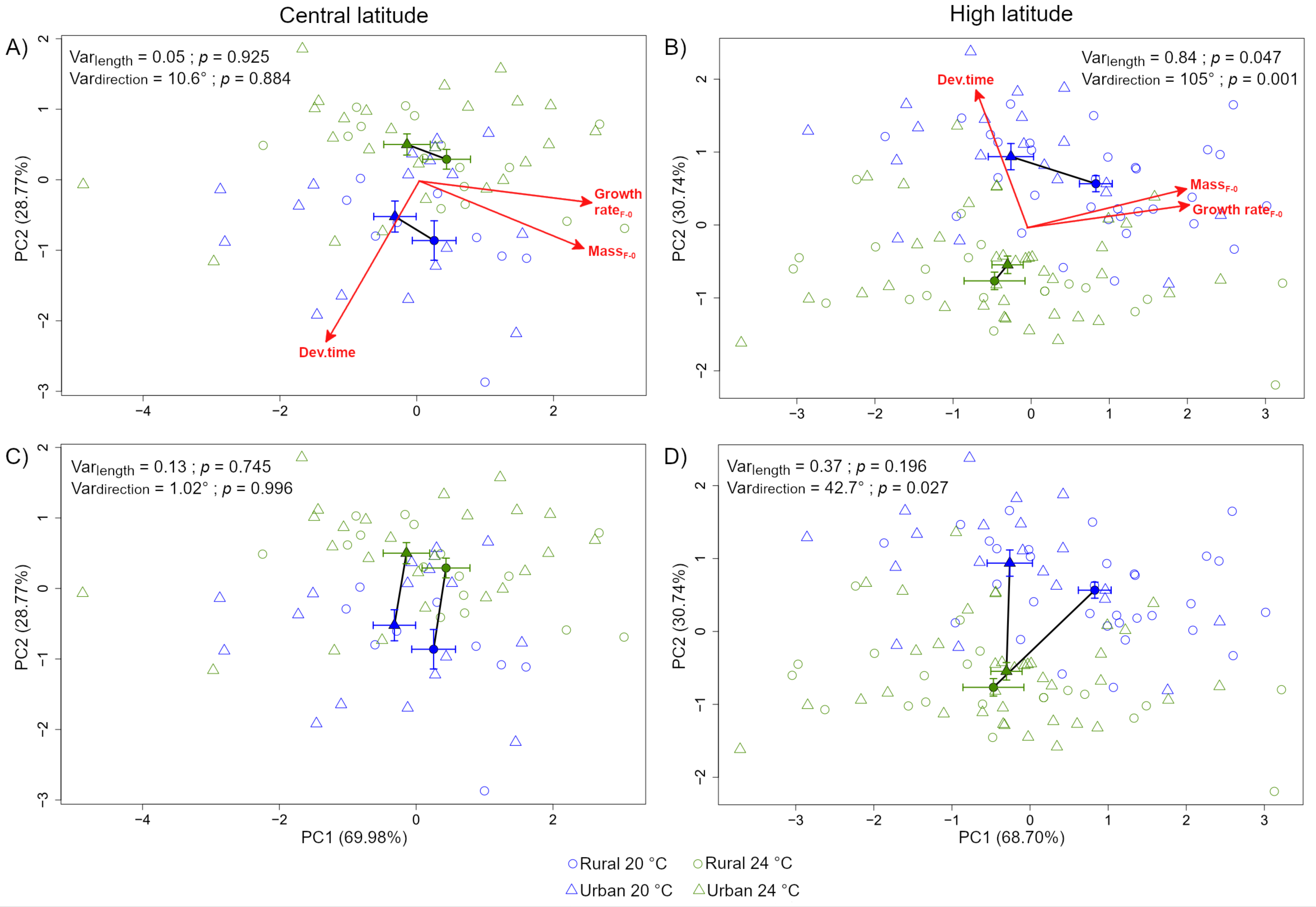

### Fig. S7.pdf

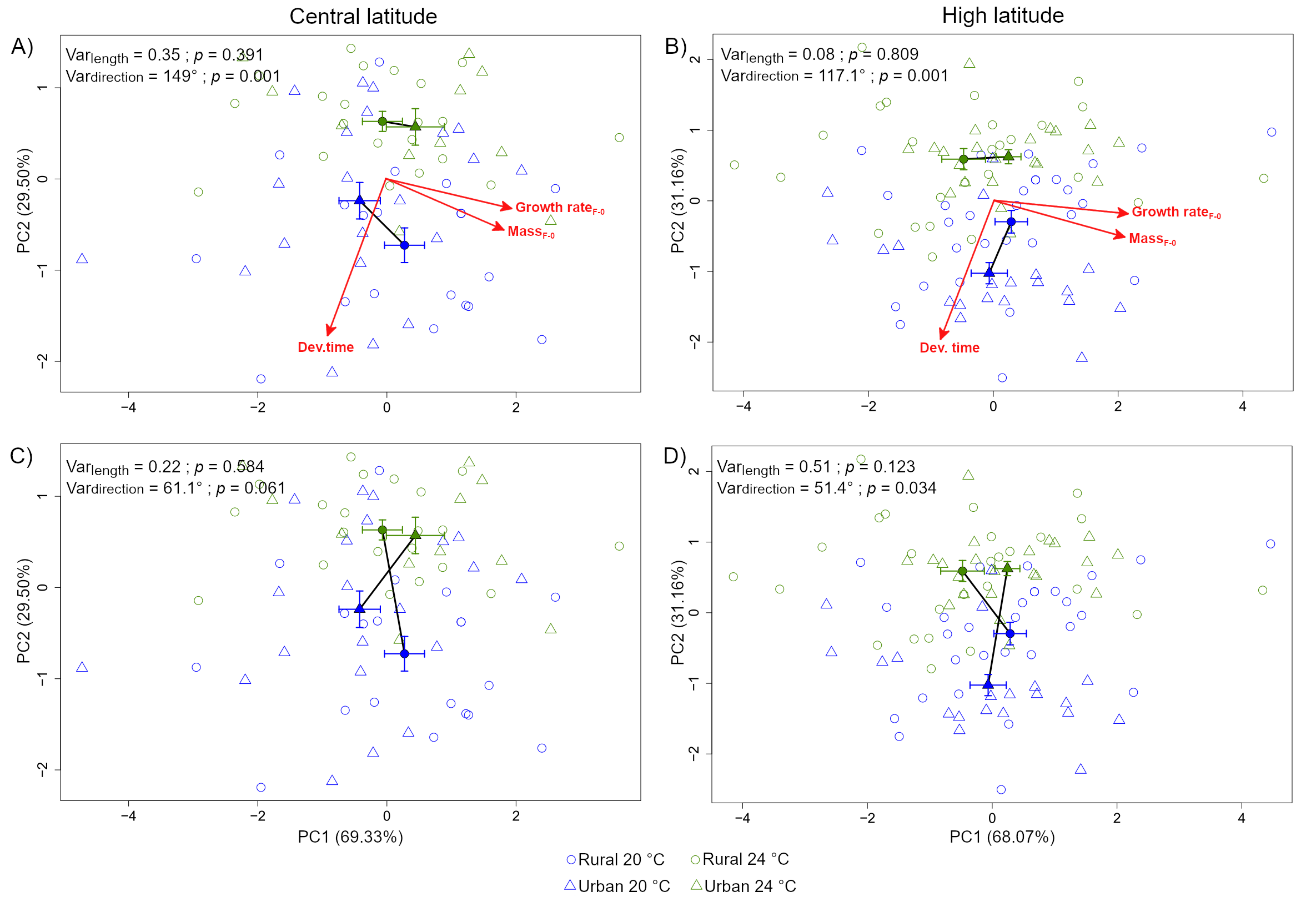

### Fig. S9.pdf

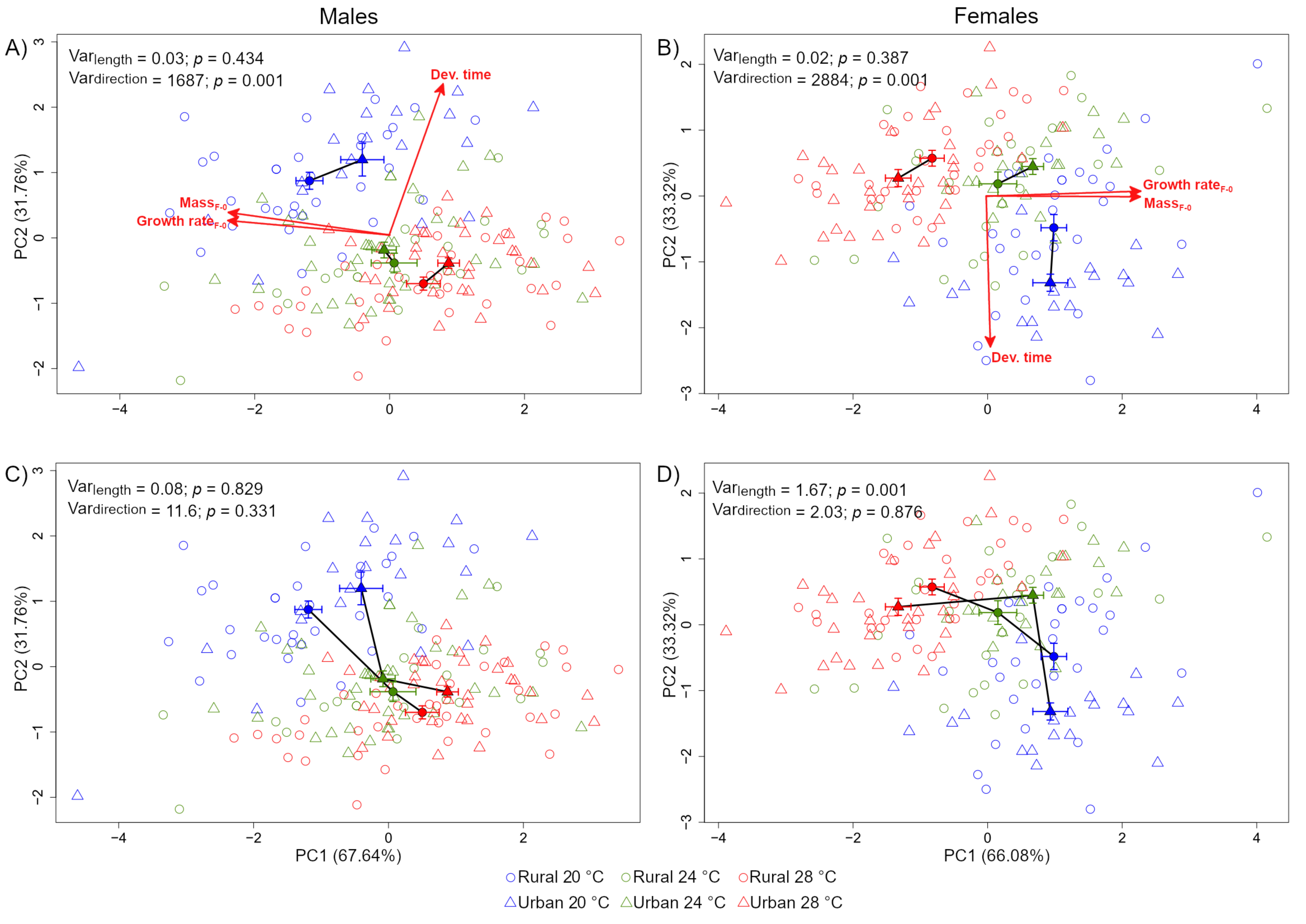
