## Supplementary material for "Latitude-specific urbanization effects on life history traits in the damselfly *Ischnura elegans*": Fig S2-S6; Fig. S8; Table S1;S2 and S5: Supp info_20_02_2023.docx

**Electronic Supplementary Material**

Guillaume Wos

Szymon Sniegula

Separate files

**Fig. S1**. Monthly temperatures for each Swedish (SW; high latitude) and Polish (PL; central latitude) ponds across urban and rural landscapes depicted by the type of lines (solid = rural; broken = urban). The figure shows temperatures for (A) central- and (B) high latitude ponds estimated using Flake (Lake Model Flake 2009) over the period 1998 – 2009. As FLake model does not consider impervious surface as a parameter. Dataloggers were installed (50 cm depth) in each pond for one year or only several months spanning the summer period in (C) central- and (D) high latitude ponds. For the urban pond located in Krakow (panel C), we placed two dataloggers, one exposed to the sun “Krakow (PL) - sun” and one in shady area “Krakow (PL) - shade” to catch the variability of temperatures within a single pond.

**Fig S6**. Principal component analysis (PCA) conducted on males only and showing evolutionary changes in response to urbanization at current (20 °C) and mild warming (24 °C) temperature for (A) central- and (B) high-latitude populations and plastic changes in response to different temperatures in rural and urban populations for (C) central- and (D) high-latitude populations. The PCAs were run on response variables measured at the larval entrance into the final instar, and before the treatment with predator cue. Rural and urban individuals are depicted by open circles and triangles, respectively; temperature by colours, blue = 20 °C and green = 24 °C; filled circles and triangles correspond to the centroid of each group; solid lines connecting filled symbols represent the vector.

**Fig S7**. Principal component analysis (PCA) conducted on females only and showing evolutionary changes in response to urbanization at current (20 °C) and mild warming (24 °C) temperature for (A) central- and (B) high-latitude populations and plastic changes in response to different temperatures in rural and urban populations for (C) central- and (D) high-latitude populations. The PCAs were run on response variables measured at the larval entrance into the final instar, and before the treatment with predator cue. Rural and urban individuals are depicted by open circles and triangles, respectively; temperature by colours, blue = 20 °C and green = 24 °C; filled circles and triangles correspond to the centroid of each group; solid lines connecting filled symbols represent the vector.

**Fig S9**. Principal component analysis (PCA) showing evolutionary changes in response to urbanization at current (20 °C), mild warming temperature (24 °C) and heat wave (28 °C) for (A) males and (B) females and plastic changes in response to temperature in rural and urban populations for (C) males and (D) females from high-latitude populations. The PCAs were run on response variables measured at the larval entrance into the final instar, and before the treatment with predator cue. Rural and urban individuals are depicted by open circles and triangles respectively; temperature by colours (blue = 20 °C, green = 24 °C and red = 28 °C); filled circles and triangles correspond to the centroid of each group; solid lines connecting filled symbols represent the vector.

**Fig. S2**. Details on (A) rearing temperatures and (B) photoperiod (green line) used over the course of the experiment. Panel A indicates the wintering period (grey area; 6 ºC constant). The end of each curve to the right indicates the end of the experiment, i.e. when last F-0 larva ended the five-day-long predator cue treatment.

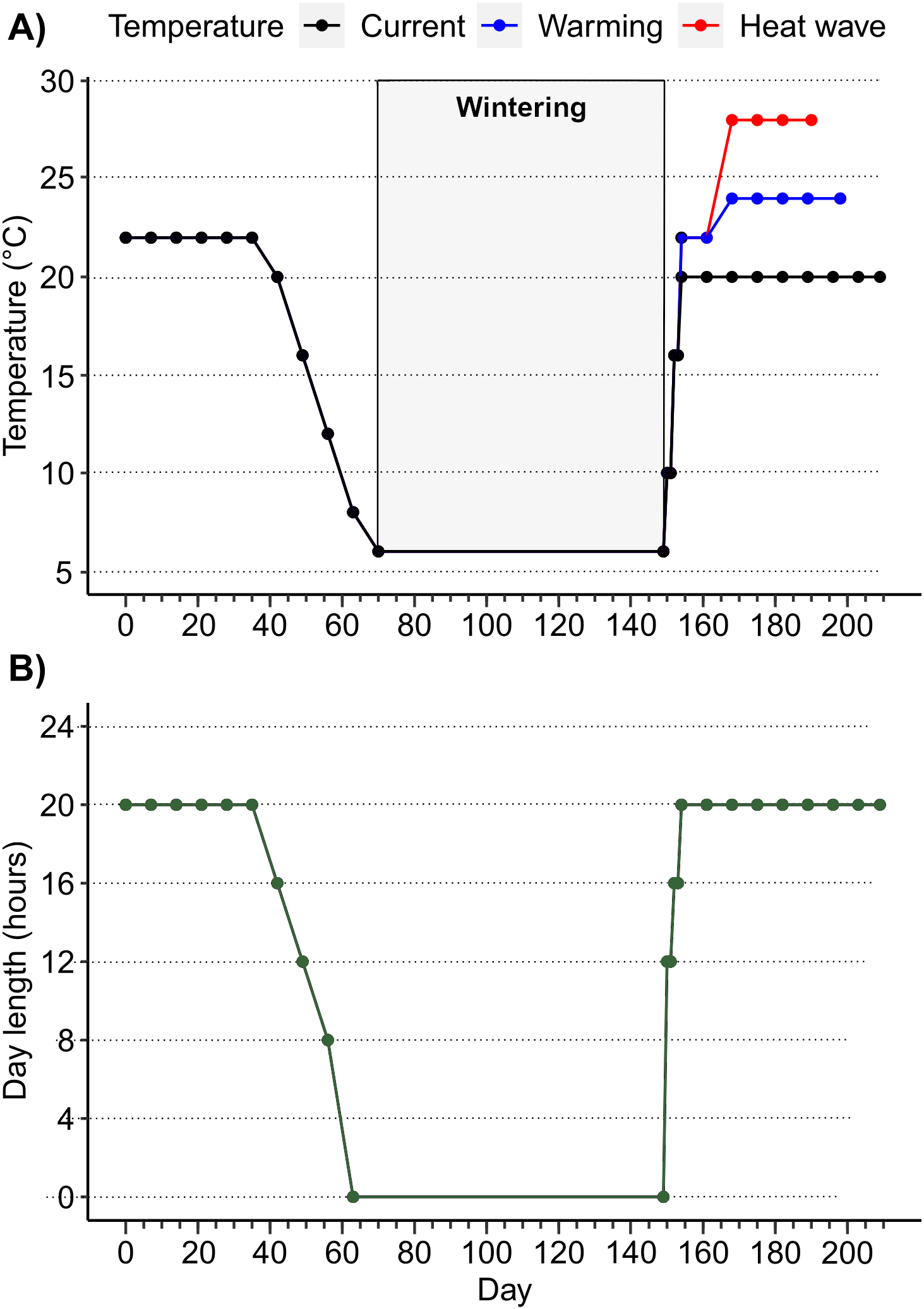

**Fig. S3**. Larval (A) mass _F0_ of females and males and (B) growth rate _F0_ in 20 °C and 24 °C, for central- and high latitude populations in Experiment 1.

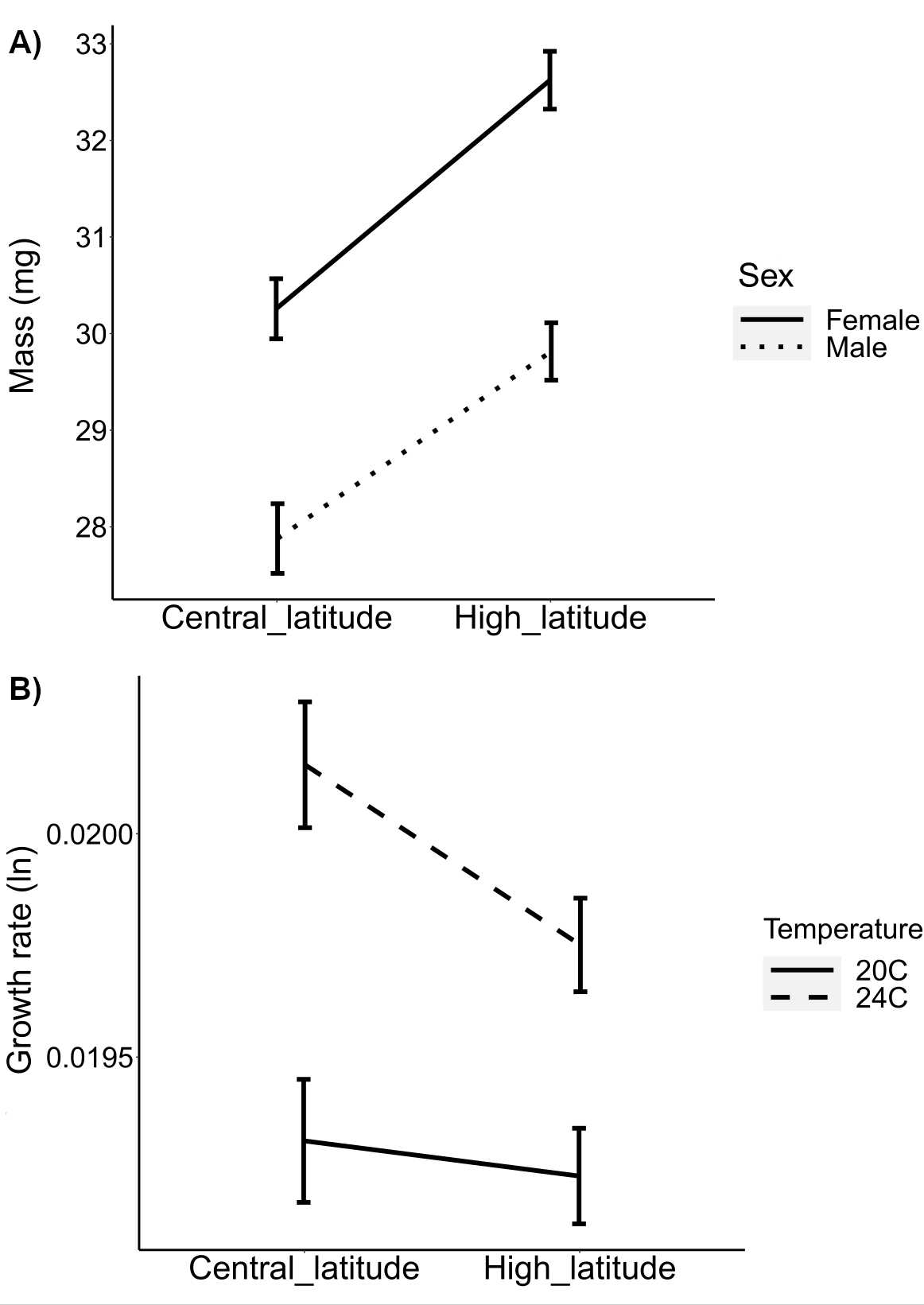

**Fig. S4.** Larval development time (in days) for central- and high latitude populations in different temperature treatments. Black horizontal lines represent the median and the box the 25th and 75th percentiles. Vertical bars are for standard errors and outliers are represented by dots.

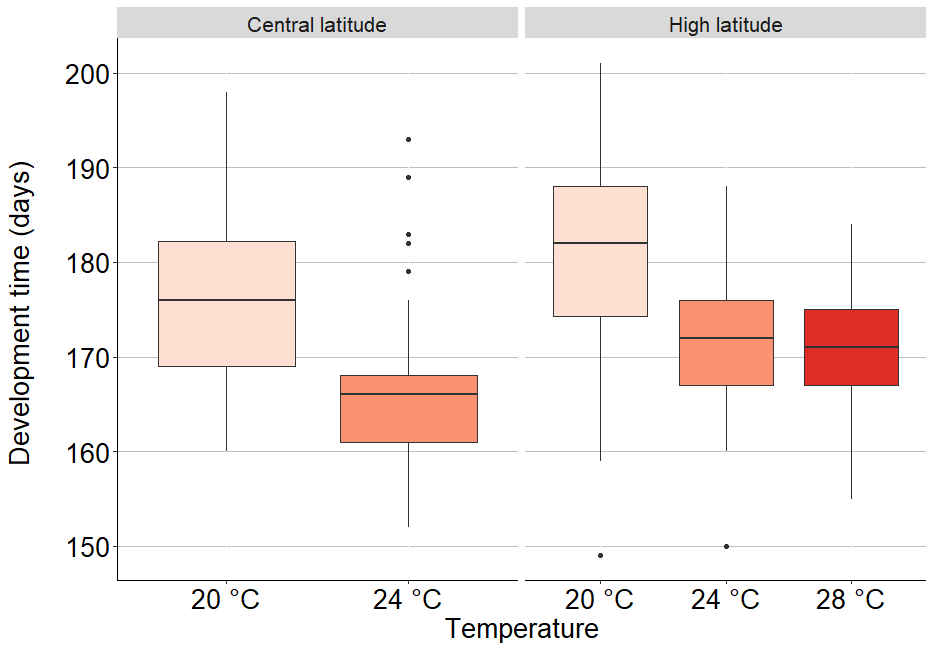

**Fig. S5.** Growth rate _final_ (ln) for A) central- and high-latitude populations and B) each predator treatment (absence vs. presence of a predator cue) for Experiment 1. Black horizontal lines represent the median and the box the 25th and 75th percentiles. Vertical bars are for standard errors and outliers are represented by dots.

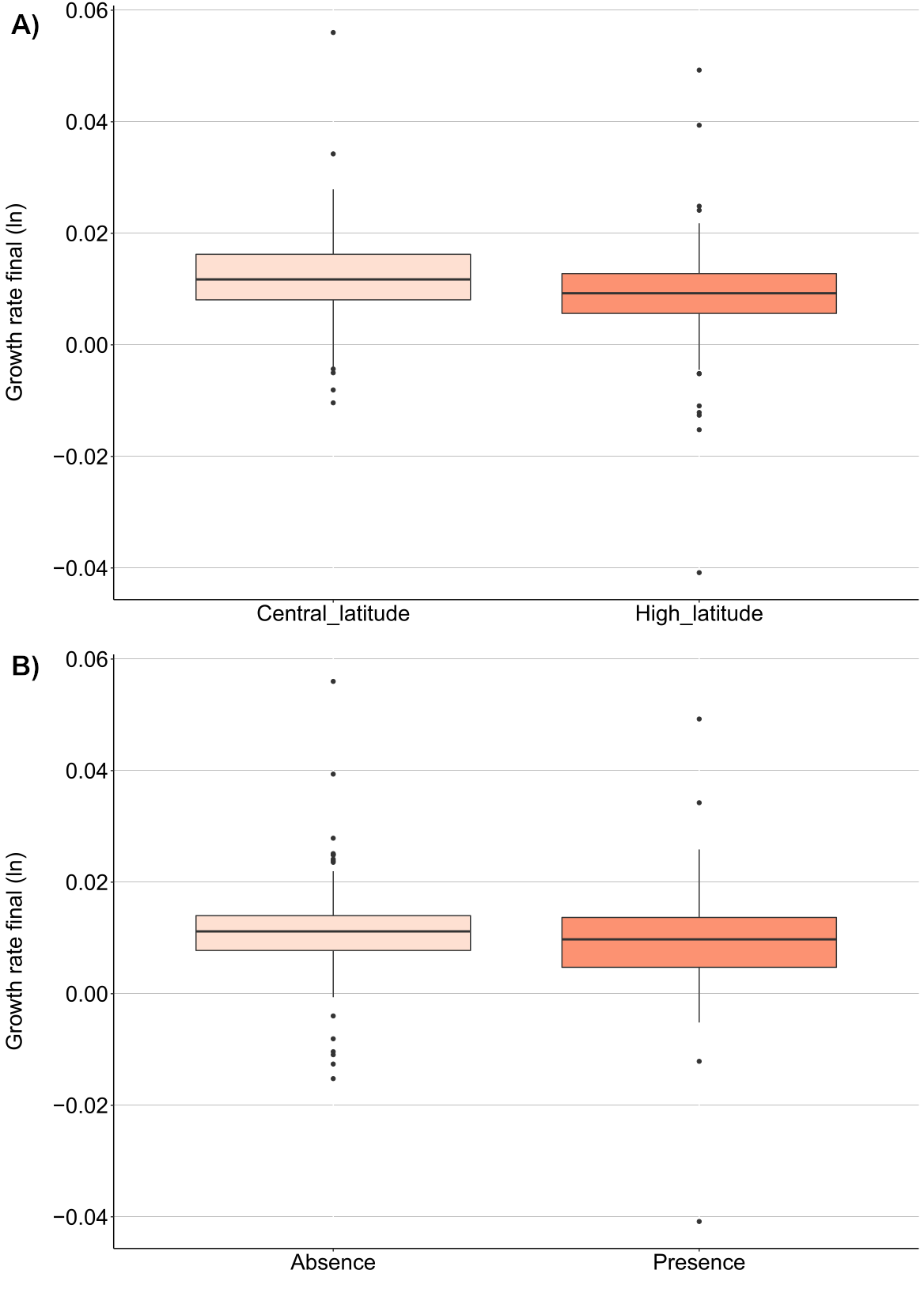

**Fig. S8**. Larval (A) mass _F0_ and (B) growth rate _F0_ (GR _F-0_) at current (20 °C), mild warming (24 °C) and heat wave (28 °C) temperature for females and males in Experiment 2.

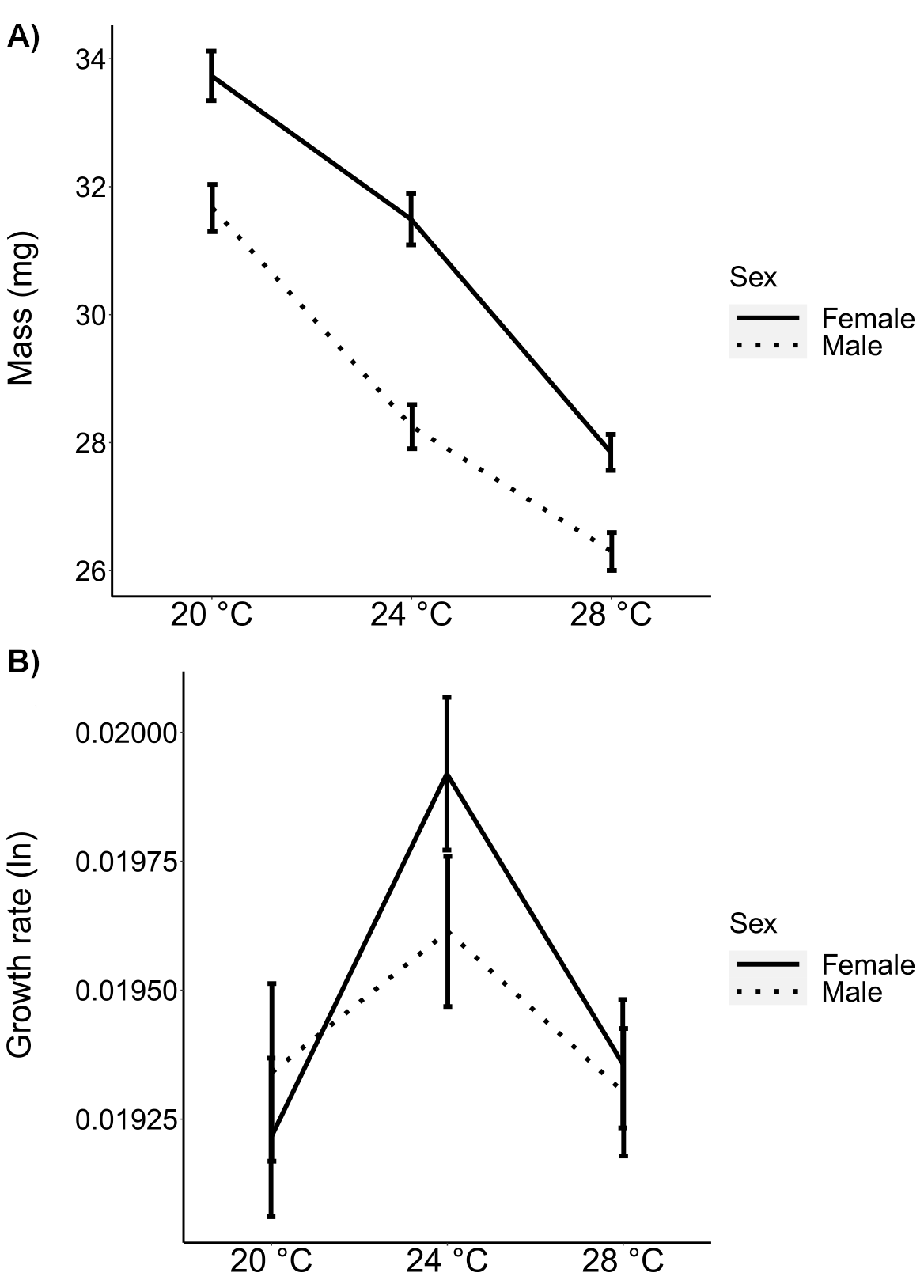

**Table S1**. Description of the sampled ponds.

|  |  |  |  | **Urbanization** | | **GPS coordinates** | |
| --- | --- | --- | --- | --- | --- | --- | --- |
| **Locality** | **Country** | **Latitude** | **Elevation (m)** | **Type** | **% impervious surface** | **Latitude** | **Longitude** |
| Torups | Sweden | High | 32.0 | Rural | 1.35 | 55.564293 | 13.205614 |
| Vallkärra | Sweden | High | 24.0 | Rural | 0.19 | 55.738166 | 13.153274 |
| Kronoborg | Sweden | High | 4.0 | Urban | 38.15 | 55.591598 | 12.993968 |
| Lund | Sweden | High | 24.0 | Urban | 25.07 | 55.705091 | 13.203357 |
| Zagorze | Poland | Central | 275.0 | Rural | 0.37 | 50.083352 | 19.39736 |
| Niepolomice | Poland | Central | 192.0 | Rural | 0.00 | 50.10875 | 20.348707 |
| Krakow | Poland | Central | 201.0 | Urban | 24.19 | 50.064616 | 19.986129 |
| Katowice | Poland | Central | 279.0 | Urban | 32.76 | 50.280691 | 19.021941 |

**Table S2**. Sample size (*N*) per explanatory variable in Experiment 1 and Experiment 2 at the start of post-winter treatments.

|  |  | **By variable** | | | | | | | | | | | |
| --- | --- | --- | --- | --- | --- | --- | --- | --- | --- | --- | --- | --- | --- |
| **Experiment 1** | **Total** | **Latitude** | | **Urbanization** | | **Sex** | | **Temperature** | | | | **Predator cue** | |
|  |  | Central | High | Rural | Urban | Male | Female | 20 ºC | | 24 ºC | | Presence | Absence |
| *N* | 349 | 141 | 208 | 174 | 175 | 179 | 170 | 169 | | 180 | | 179 | 170 |
| **Experiment 2** | **Total** | **Latitude** | | **Urbanization** | | **Sex** | | **Temperature** | | | | **Predator cue** | |
|  |  | High | | Rural | Urban | Male | Female | 20 ºC | 24 ºC | | 28 ºC | Presence | Absence |
| *N* | 340 | 340 | | 173 | 167 | 182 | 158 | 101 | 107 | | 132 | 174 | 166 |

**Table S5**. Least square mean values and standard errors for each response variable in Experiment 1 and Experiment 2.

| Experiment 1 |  | Entrance into the final instar F-0 | | | Predator treatment |
| --- | --- | --- | --- | --- | --- |
|  |  | Dev. time (days) | Mass (mg) | Growth rate (ln) | Growth rate (ln) |
| **Sex** | **Male** | 173 ± 0.73 | 29.1 ± 0.23 | 19.5e-3 ± 8.7e-5 | 10.6e-3 ± 6.1e-4 |
|  | **Female** | 175 ± 0.73 | 31.6 ± 0.24 | 19.7e-3 ± 8.9e-5 | 9.8e-3 ± 6.2e-4 |
| **Latitude** | **Central** | 171 ± 0.80 | 29.1 ± 0.27 | 19.7e-3 ± 9.8e-5 | 12.2e-3 ± 6.8e-4 |
|  | **High** | 176 ± 0.66 | 31.1 ± 0.22 | 19.5e-3 ± 8.0e-5 | 8.8e-3 ± 5.6e-4 |
| **Temperature** | **20 ˚C** | 179 ± 0.65 | 31.6 ± 0.24 | 19.3e-3 ± 8.6e-5 | 9.7e-3 ± 6.3e-4 |
|  | **24 ˚C** | 169 ± 0.63 | 29.1 ± 0.23 | 19.9e-3 ± 8.4e-5 | 10.8e-3 ± 6.1e-4 |
| **Urbanization** | **Rural** | 174 ± 0.74 | 30.8 ± 0.25 | 19.7e-3 ± 8.8e-5 | 10.6e-3 ± 6.2e-4 |
|  | **Urban** | 174 ± 0.74 | 29.8 ± 0.25 | 19.5e-3 ± 8.8e-5 | 9.8e-3 ± 6.2e-4 |
| **Predator** | **Absence** | / | / | / | 11.2e-3 ± 6.1e-4 |
|  | **Presence** | / | / | / | 9.1e-3 ± 6.2e-4 |

| Experiment 2 |  | Entrance into the final instar F-0 | | | Predator treatment |
| --- | --- | --- | --- | --- | --- |
|  |  | Dev. time (days) | Mass (mg) | Growth rate (ln) | Growth rate (ln) |
| **Sex** | **Male** | 173 ± 0.63 | 28.5 ± 0.26 | 19.4e-3 ± 8.1e-5 | 9.5e-3 ± 5.9e-4 |
|  | **Female** | 176 ± 0.67 | 30.8 ± 0.28 | 19.5e-3 ± 8.7e-5 | 7.8e-3 ± 5.5e-4 |
| **Temperature** | **20 ˚C** | 181 ± 0.72 | 32.7 ± 0.28 | 19.3e-3 ± 10.7e-5 | 8.7e-3 ± 7.4e-4 |
|  | **24 ˚C** | 172 ± 0.71 | 29.7 ± 0.27 | 19.8e-3 ± 10.4e-5 | 8.9e-3 ± 7.2e-4 |
|  | **28 ˚C** | 171 ± 0.63 | 27.0 ± 0.25 | 19.3e-3 ± 9.4e-5 | 8.5e-3 ± 6.5e-4 |
| **Urbanization** | **Rural** | 173 ± 0.65 | 29.8 ± 0.28 | 19.6e-3 ± 8.3e-5 | 8.6e-3 ± 5.7e-4 |
|  | **Urban** | 175 ± 0.66 | 29.3 ± 0.28 | 19.3e-3 ± 8.4e-5 | 8.7e-3 ± 5.8e-4 |
| **Predator** | **Absence** | / | / | / | 8.8e-3 ± 5.7e-4 |
|  | **Presence** | / | / | / | 8.6e-3 ± 5.8e-4 |
